# Single-cell RNA-seq analysis reveals penaeid shrimp hemocyte subpopulations and cell differentiation process

**DOI:** 10.1101/2021.01.10.426076

**Authors:** Keiichiro Koiwai, Takashi Koyama, Soichiro Tsuda, Atsushi Toyoda, Kiyoshi Kikuchi, Hiroaki Suzuki, Ryuji Kawano

## Abstract

Crustacean aquaculture is expected to be a major source of fishery commodities in the near future. Hemocytes are key players of the immune system in shrimps; however, their classification, maturation, and differentiation are still under debate. To date, only discrete and inconsistent information on the classification of shrimp hemocytes has been reported, showing that the morphological characteristics are not sufficient to resolve their actual roles. Our present study using single-cell RNA sequencing, revealed nine types of hemocytes of *Marsupenaeus japonicus* based on their transcriptional profiles. We identified markers of each subpopulation and the differentiation pathways involved in their maturation. We also discovered cell growth factors that might play crucial roles in hemocyte differentiation. Different immune roles among these subpopulations were suggested from the analysis of differentially expressed immune-related genes. These results provide a unified classification of shrimp hemocytes, which improves the understanding of its immune system.

## Introduction

Aquaculture is an important source of animal protein and is considered one of the most important long-term growth areas of food production, providing 60% of fish for human consumption (1) (http://www.fao.org/fishery/statistics/en). However, crustaceans that lack an adaptive immune system (2-4) are vulnerable to pathogens. This means that ordinal vaccination is not applicable to crustaceans, unlike in fish aquaculture. Shrimp is the main target species for crustacean aquaculture. Therefore, an immune priming system for shrimp, which is entirely different from conventional vaccines, needs to be developed to control the infection of pathogens. However, little is known about the immune system of crustaceans due to the lack of biotechnological tools, such as uniform antibodies and other biomarkers (5).

Hemocytes, which are immune cells of crustaceans, are traditionally divided into three morphological types based on the dyeing of intracellular granules, which was established by Bauchau and colleagues (6-8). However, there have been additional reports on the classification of the hemocytes of shrimp; they were classified into four, eight, and five types based on electron microscopy (9), another dyeing method (10), and iodixanol density gradient centrifugation (11), respectively. As the morphology and dye staining properties of shrimp hemocytes are not absolute indicators, no unified understanding of their role has been established yet. Molecular markers, such as specific mRNAs, antibodies, or lectins, are usually available for characterizing the subpopulations of cells in model organisms, but this is not often the case for non-model organisms. Although monoclonal antibodies have been developed for some hemocytes of shrimp (12-18), their number is lower than that of humans, and their correspondence to the cell type, as well as their differentiation stage are under debate.

Recently, single-cell mRNA sequencing (scRNA-seq) techniques have dramatically changed this scene, allowing researchers to annotate non-classified cells solely based on the mRNA expression patterns of each cell. In particular, droplet-based mRNA sequencing, such as Drop-seq, developed by Macosko, Basu (19), has gained popularity for classifying cells and identifying new cell types. The enormous amount of biological data obtained from scRNA-seq leads us to classify cells into specific groups, analyze their heterogeneity, predict the functions of single-cell populations based on the gene expression profiles, and determine the cell proliferation or development pathways based on the pseudo-time ordering of a single cell (20, 21). More recently, hemocytes of invertebrate, fly, and mosquito models have been subjected to these types of microfluidic-based scRNA-seq to reveal their functions (22, 23).

Here, we performed scRAN-seq analysis on *Marsupenaeus japonicus* hemocytes to classify the hemocyte types and to characterize their functions using the custom-built Drop-seq platform. To perform scRNA-seq, a high-quality gene reference is essential; however, such reference genomes are scarce for crustaceans because of the extremely high proportion of simple sequence repeats (5). We circumvented this problem by preparing reference genomes using hybrid *de novo* assembly of short- and long-read RNA sequencing results. The sequences obtained from the scRNA-seq were mapped onto the reference genes successfully. Our scRNA-seq uncovered the transcriptional profiles of a few thousand *M. japonicus* hemocytes. We identified the markers of each population and the differentiation pathways associated with their maturation. We also discovered the cell growth factors that might play crucial roles in hemocyte differentiation. Different immune roles among these subpopulations were also suggested from the analysis of differentially expressed immune-related genes. Our results present a unified classification of shrimp hemocytes and a deeper understanding of the immune system of shrimp.

## Results

### scRNA-seq clustering of *Marsupenaeus japonicus* hemocytes

Our study utilized scRNA-seq to determine the cellular subtypes with a distinct transcriptional expression (Fig. 1 A). To map the scRNA-seq sequences from *M. japonicus* hemocytes, we first prepared *de novo* assembly of the reference genes using hybrid assembly of short- and long-read RNA sequencing results. Then, by using self-built Drop-seq microfluidic chips, single hemocytes were captured and their mRNA was barcoded using the droplet-based strategy. This process was performed in triplicates for three shrimp individuals. Following library preparation and sequencing, the transcriptomes obtained from scRNA-seq were mapped against the reference genes to discover the cell types.

**Figure 1.**
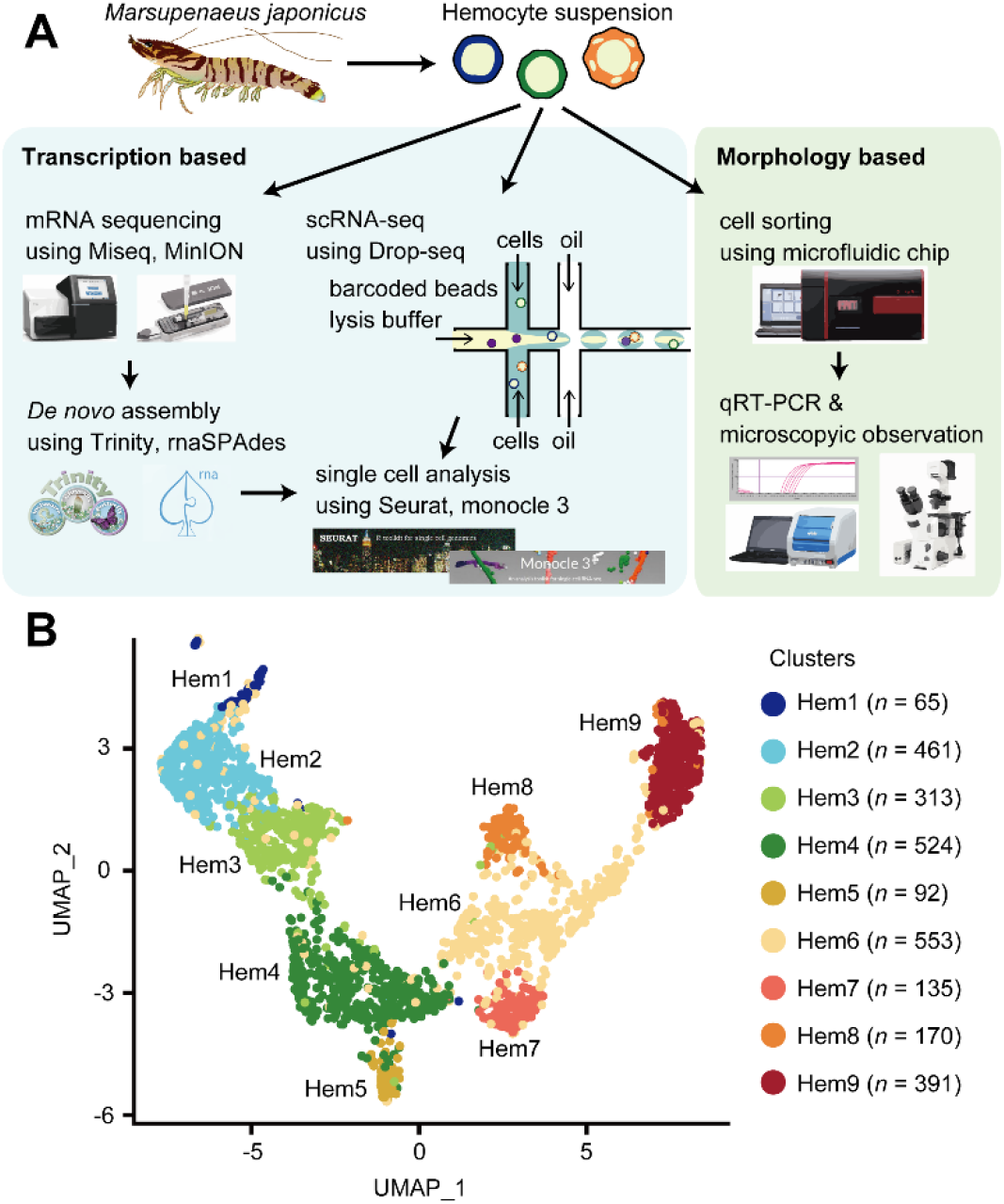
scRNA-seq of penaeid shrimp *M. japonicus* hemocytes. (A) Schematics of the microfluidics-based scRNA-seq, preparation of the *de novo* assembled gene list, *in silico* analysis workflow and morphology-based cell classification. (B) UMAP (uniform manifold approximation and projection) plot of SCTransform batch corrected and integrated of hemocytes from three shrimps (*n* = 2,704).

Using the Drop-seq procedure, we profiled a total of 2,704 cells and obtained a median value of 718 unique molecular identifiers (UMIs) and 334 genes per cell across three replicates (Fig. S1 A and B). Approximately 300 genes were detected, and the total number of mRNAs expressed among individual cells varied between 100 and 1,400 (Fig. S1 C and D). There was some transcriptional variability, which may have resulted from the artifacts of the Drop-seq system, because it is consistent with the original Drop-seq paper (19) and with recent findings in other organisms, such as fish (24), flies (23), and mosquitoes (22, 25).

Applying the SCTransform batch correction method integrated into the Seurat package allowed us to remove the individual differences. SCTransform successfully integrated all three shrimp datasets, among which we identified a total of nine clusters (Fig. 1 B) and obtained 3,334 commonly expressed genes. Each cluster contained the following number of cells: Hem 1, 65 cells (2.4%); Hem 2, 461 cells (17.0%); Hem 3, 313 cells (11.6%); Hem 4, 524 cells (19.4%); Hem 5, 92 cells (3.4%); Hem 6, 553 cells (20.5%); Hem 7, 135 cells (5.0%); Hem 8, 170 cells (6.3%); and Hem 9, 391 cells (14.5%), respectively.

### Cluster specific markers and their functional prediction

A total of 40 cluster-specific markers were predicted using the Seurat FindMarkers tool (Fig. 2, Dataset S1, and Dataset S2). For each cluster, seven (Hem 1), six (Hem 2), one (Hem 3), four (Hem 4), eight (Hem 5), five (Hem 7), one (Hem 8), and eight (Hem 9) markers were selected. Their functions were then annotated using BLASTX searching for penaeid shrimp identical proteins downloaded from a public database.

**Figure 2.**
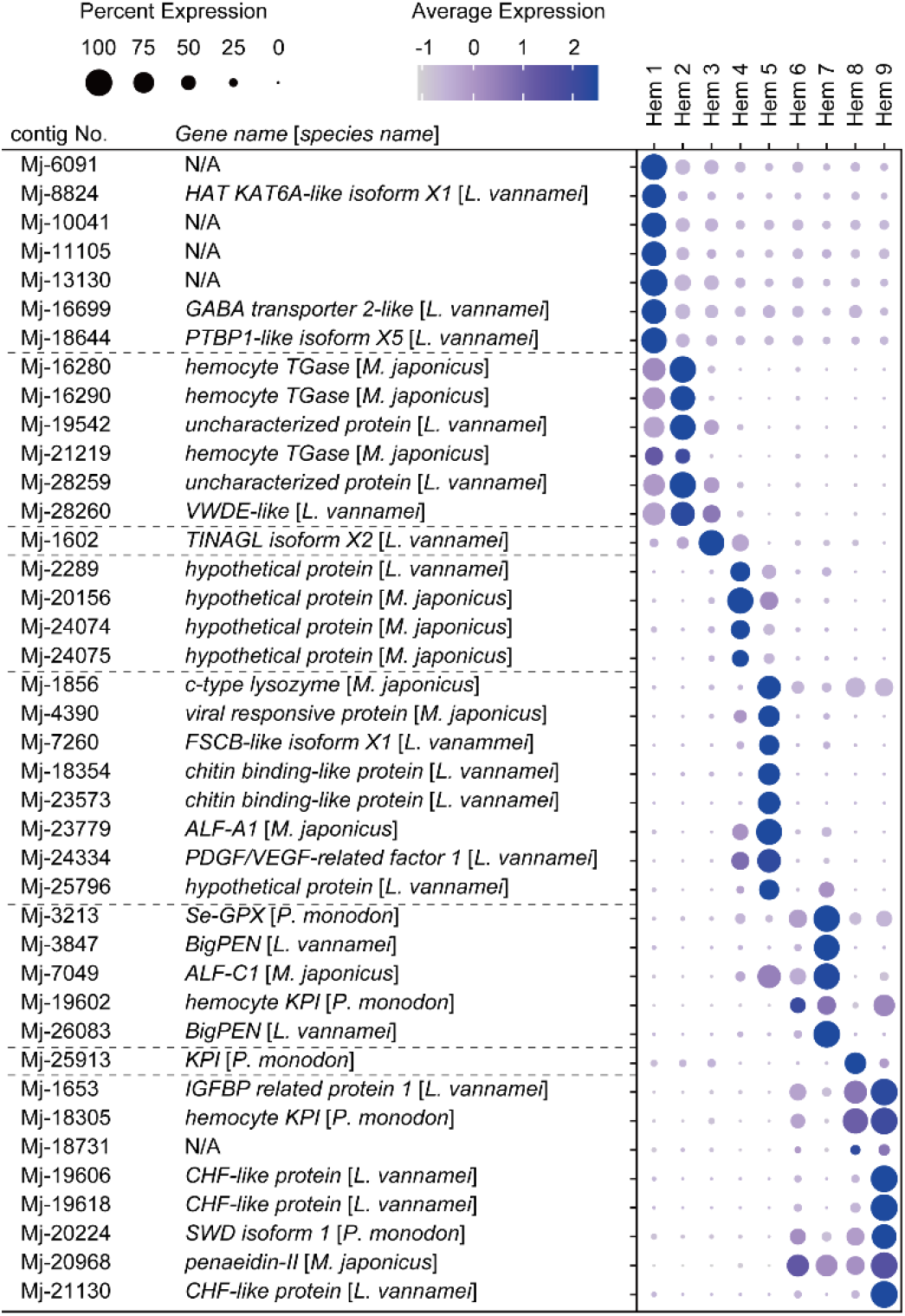
Dot plot representing the marker genes per cluster based on the average expression predicted using the Seurat FindMarker tool. Color gradient of the dot represents the expression level, while the size represents the percentage of cells expressing any gene per cluster.

Hem 1 specific markers, *histone acetyltransferase* lysine acetyltransferase 6A (*HAT KAT6A), gamma-aminobutyric acid* (*GABA*) *transporter*, and *polypyrimidine tract-binding protein 1* (*PTBP1*) are genes related to cell proliferation, cell migration, and colony formation in human tumor studies. HAT KAT6A is known to be a chromatin regulator that controls fundamental cellular processes and is implicated in regulating tumor progression (26). Autocrine/paracrine signaling via GABA receptors negatively controls ES cell and peripheral neural crest stem cell proliferation (27). This GABA signaling pathway critically regulates proliferation independently of differentiation, apoptosis, and overt damage to DNA (27). PTBP1 is a multi-functional RNA-binding protein that is overexpressed in glioma, a type of tumor that occurs in the brain, and a decreased expression of PTBP inhibits cell migration and increases the adhesion of cells to fibronectin and vitronectin (28, 29). PTBP has been shown to be involved in germ cell differentiation in *Drosophila melanogaster* and is essential for the development of *Xenopus laevis* (30). These pieces of evidence strongly suggest that these Hem 1 markers are related to cell proliferation in shrimp hemocytes. Among the six markers found in cluster Hem 2, three markers were annotated with hemocyte transglutaminase (HemTGase), and one marker showed high similarity with von Willebrand factor D and epidermal growth factor (EGF) domain-containing protein (VWDE). Interestingly, both Hem 1 and Hem 2 showed a high expression of TGase, an immature hemocyte marker of crayfish and shrimp. When the extracellular TGase is digested, hemocytes start to differentiate into mature hemocytes (31-33). The high expression of TGase suggests that both Hem 1 and Hem 2 are in the early stage of hemocytes. In cluster Hem 3, the only identified marker showed high similarity with tubulointerstitial nephritis antigen (TINAGL). In cluster Hem 4, all markers showed similarity with the hypothetical protein.

In clusters Hem 5 to Hem 9, many of the cluster-specific markers showed similarity with immune-related genes of penaeid shrimp, and very few genes were unknown. We postulate that this is partly because the public database is rich in immune-related genes because of their importance. In cluster Hem 5, markers showed a high similarity with c-type lysozyme, viral responsive protein (VRP), fibrous sheath CABYR-binding (FSCB)-like, chitin binding-like protein, anti-lipopolysaccharide factor (ALF)-A1, and PDGF/VEGF-related factor 1. In cluster Hem 7, markers showed high similarity with selenium-dependent glutathione peroxidase (Se-GPX), BigPEN, ALF-C1, and hemocyte Kunitz protease inhibitor (KPI). In cluster Hem 8, the marker showed high similarity with KPI. In cluster Hem 9, the markers showed similarity with insulin-like growth factor-binding protein (IGFBP)-related protein 1, hemocyte KPI, crustacean hematopoietic factor (CHF)-like protein, single whey acidic protein domain (SWD)-containing protein, and penaeidin-II.

Altogether, only 70% (28/40) of markers were annotated from the BLAST searches on the penaeid shrimp identical protein data. We also performed GO annotation using eggNOG-mapper (34, 35) (http://eggnog-mapper.embl.de/) to predict the functions of contigs, but only about 10% of the contigs were annotated, and we could not reach GO analysis.

From the results of this cluster-specific marker analysis, we found that Hem 1 and Hem 2 are immature hemocytes. Therefore, we analyzed the whole maturation process of hemocytes using pseudo-temporal analysis. Additionally, cell growth-related genes, such as *VWDE, TINGAL, IGFBP-related protein*, and *CHF*, were identified as cluster-specific markers. We predicted their functions to pursue the linkage between these genes and the hemocyte differentiation process. Finally, the immune-related genes, which were enriched in Hem 5 to 9, were analyzed to predict the immune function of each cluster.

### Pseudo temporal ordering of cells delineates hemocyte lineages

To investigate the dynamics of hemocyte differentiation, we performed lineage-tree reconstruction using the monocle3 learn_graph function. The differentiation and proliferation of hemocytes in shrimp and other crustaceans are still under debate (36). Since the cell cycle- and hemocyte-type-specific markers are better studied in *Drosophila*, we checked the commonly expressed genes of *M. japonicus* that are similar to the markers of *Drosophila* to determine whether they are present in shrimp. In the search for cell cycle-specific markers (Dataset S3), a large portion of cells in Hem 1 expressed the G2/M-related genes of *Drosophila*: *heterochromatin protein 1b* (*HP1b*); Mj-13141, *CCCTC-binding factor* (*CTCF*); Mj-14904, *suppressor of variegation 205* (*Su(var)205*); Mj-27796 (Fig. 3 A and Fig. S2). *Drosophila HP1* and *Su(var)205* are known to be essential for the maintenance of the active transcription of the euchromatic genes functionally involved in cell-cycle progression, including those required for DNA replication and mitosis (37, 38). CTCF has zinc finger domains and plays an important role in the development and cell division of fly and mammalian cells (39, 40). In the BLAST search of two genes related to cell-cycle progression, that is, Mj-13141 and Mj-27796, on identical proteins of penaeid shrimp, they showed high similarity with the *chromobox protein homolog 1-like* (Dataset S1). A chromobox family protein contributes to lymphomagenesis by enhancing stem cell self-renewal and/or by increasing the replicative potential of cancer stem cells in human tumor cells (41). BLAST search of Mj-14904 on identical proteins of penaeid shrimp showed similarity with the zinc finger protein that contains the same domain of CTCF (Dataset S1). These findings suggested that hemocytes grouped as Hem 1 are tightly regulated by these G2/M phase-related genes to promote cell division.

**Figure 3.**
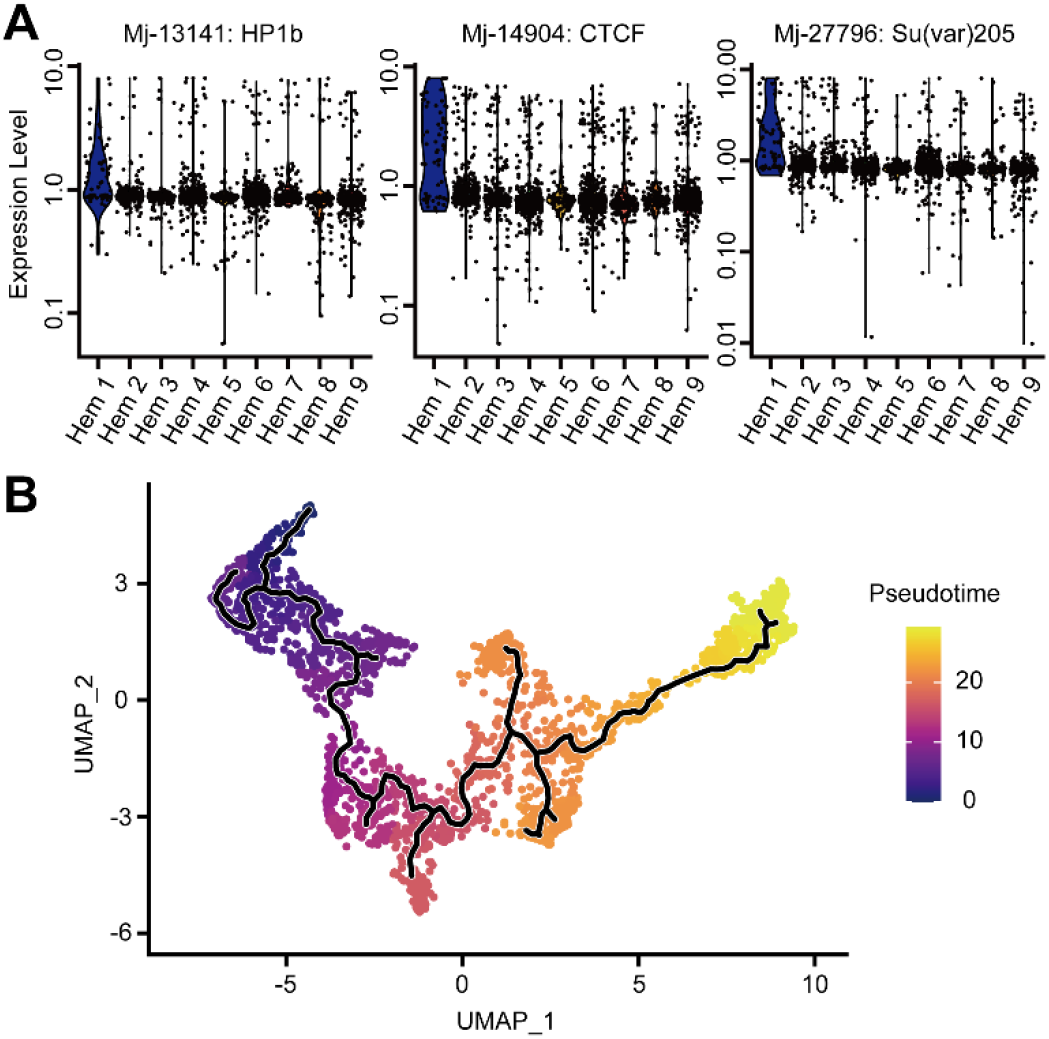
Pseudo temporal ordering of hemocyte lineages. (A) Violin plots displaying normalized expression levels of each cell cycle related genes across all clusters. (B) Visualization of clusters (from Figure 1B) onto the pseudo time map using monocle 3.

Among the specific markers of four types of *Drosophila* hemocytes, four genes in prohemocytes, 11 genes in plasmatocytes, 11 genes in lamellocytes, and 6 genes in crystal cells showed similarity with the shrimp genes (Dataset S4). However, no genes were significantly expressed in certain clusters (Fig. S3). Therefore, we concluded that these *Drosophila* hemocyte markers cannot be adapted as markers for the hemocytes of shrimp.

We considered Hem 1 to be the initial state of hemocytes and set it as the starting point in the differentiation process because Hem 1 expressed cell proliferation-related genes, TGase (an immature hemocyte marker), and G2/M phase-related genes (37-42). From this pseudo temporal ordering analysis, we found four main lineages starting from Hem 1 to Hem 5, Hem 7, Hem 8, and Hem 9 at the endpoints (Fig. 3 B and Fig. S4 A-D). In crayfish, hematopoietic stem cells are present in hematopoietic tissue (HPT), and two types of hemocyte lineages starting from a hematopoietic stem cell exist (43). In *Penaeus monodon*, hyaline cells (i.e., agranulocytes) are considered as the young and immature hemocytes of two types of matured hemocytes (9). Our pseudo temporal ordering analysis revealed that the hemocytes of *M. japonicus* differentiate from a single subpopulation into four major populations. The differentiation process of *M. japonicus* hemocytes was continuous, not discrete, which was in agreement with previous arguments on the crustacean hematopoiesis mechanism (9, 43, 44).

### Expression of cell growth-related genes

The exploration of cluster-specific markers revealed that cell growth-related genes were specifically expressed in certain clusters. Therefore, we highlighted five genes that are predicted to be involved in cell growth and differentiation: *VWDE-like, TINAGL, PDGF/VEGF-related factor 1, IGFBP-related protein 1*, and *CHF-like* (Fig. 4, Fig. S5, and Dataset S5).

**Figure 4.**
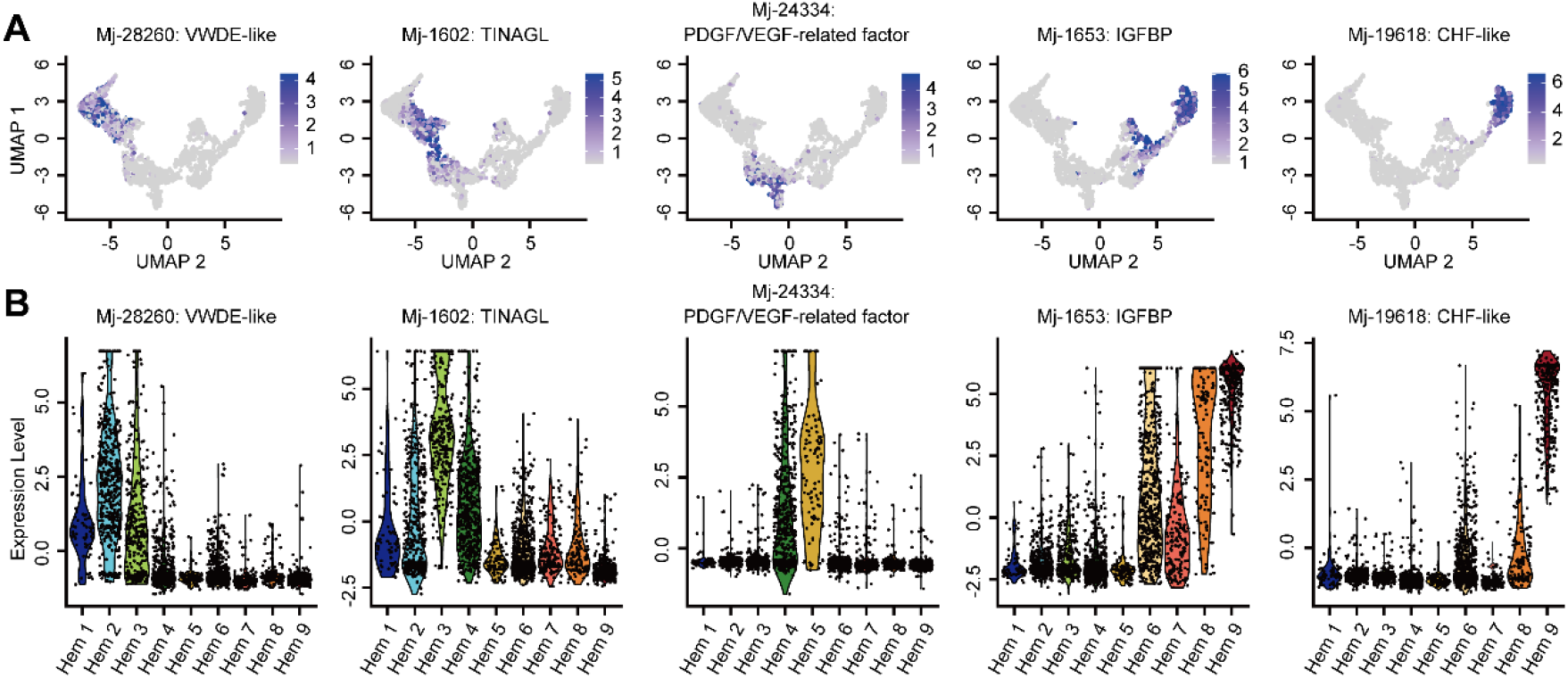
Cell growth-related gene expression across all clusters. (A) Expression pattern of the cell growth-related genes across cell clusters. (B) Violin plots displaying normalized expression levels of each cell growth-related genes across all clusters.

VWDE-like was highly expressed in clusters Hem 1 to Hem 3 (Fig. 4 A and B). The expression of VWDE is a common feature of blastemas, which are capable of regenerating limbs and fins in a variety of highly regenerative species (45). VWDE contains several epidermal growth factor (EGF) domains and is expected to be a downstream effector once a blastema has been established, but it is not a driver of blastema formation (45). VWDE-like of *M. japonicus* was expressed more strongly in Hem 2 than in Hem 1, indicating that VWDE-like also works as a downstream effector against Hem 1, which is predicted to be composed of undifferentiated hemocytes.

TINAGL, which was expressed in Hem 3 and 4, is a secreted extracellular protein that is essential for early angiogenesis in developing zebrafish embryos (46) and humans (47). TINAGL of *M. japonicus* has been proposed to participate in angiogenic cell differentiation. TINAGL is also known as a suppressor of cancer progression and metastasis by binding to receptors of EGF in humans (48). Interestingly, TINAGL of *M. japonicus* was expressed in Hem 3 and Hem 4, in which the expression of VWDE-like was weakened (Fig. 4 A and B). This suggests that TINAGL might suppress the differentiation function of VWDE-like in Hem 3 and 4 in shrimp.

Cells expressing PDGF/VEGF-related factors were dominant in Hem 4 and Hem 5 (Fig. 4 A and B). The vascular endothelial growth factor (VEGF) signaling pathway is essential for vasculogenesis, cell proliferation, and tumor migration in mammals (49, 50). Furthermore, in *Drosophila*, VEGF homologs control the number of circulating hemocytes (51). The high expression of PDGF/VEGF-related factors in Hem 4 and Hem 5 showed that Hem 3 would differentiate into Hem 5 through Hem 4 via the VEGF signaling pathway.

Hem 6 to Hem 9 expressed insulin-like growth factor (IGF) binding protein (IGFBP) (Fig. 4 A and B). IGFBP delivers IGFs to the target cells in mammal studies and is essential for cell growth or differentiation (52). The high expression of a receptor of the insulin-like peptide at mature hemocytes in the mosquito suggests that the insulin signaling pathway regulates hemocyte proliferation (53). The silencing and overexpression of IGFBP caused a decrease and increase in the growth of hemocytes, respectively, in the loss and gain function study of abalone *Haliotis diversicolor* (54). These studies indicate that the IGFBP-related insulin signaling pathway is important for hemocyte proliferation and differentiation in invertebrates. IGFBP might play an essential role in the differentiation of hemocytes from Hem 4 to Hem 6–9 in shrimp. The high expression of IGFBP is determined in the brain and gonads of *Litopenaeus vannamei* (55). This fact also suggests that IGFBP plays a possible role in organ growth and maturation in shrimp.

Up to this point, the analyzed genes were proliferation- and differentiation-promoting, but there was also the specific expression of the hemocyte homeostasis regulatory gene, crustacean hematopoietic factor (CHF) (43) (Fig. 4 A and B). CHF is a hematopoietic factor of crayfish, and the silencing of CHF leads to an increase in the apoptosis of cells in HPT, and a reduction in the number of circulating hemocytes (33). Additionally, the silencing of laminin, a receptor of CHF, reduces the number of circulating hemocytes by decreasing the number of agranulocytes, as opposed to granulocytes, in *P. vannamei* (55). CHF-like was expressed at cluster Hem 9 (Fig. 4 A and B), in which hemocytes are predicted to be matched here. Taken together, CHF-like expressed from matured hemocytes, Hem 9, might work as a hematopoietic factor against agranulocytes or regulate the homeostasis of agranulocytes.

### Expression of immune-related genes in single cells

Hemocytes of shrimp play key roles in their immunity; therefore, we selected immune-related genes from 3,334 commonly expressed genes, and then analyzed their expression levels to deduce the detailed immune functions of each cluster.

Antimicrobial peptides (AMPs) play the most important role in the immunity of shrimps and are well known to be stored in granulocytes (56, 57). The expression patterns of AMPs revealed that the major AMPs of penaeid shrimp were expressed in clusters Hem 5 to Hem 9 (Fig. 5 A-F, Fig. S6 and Dataset S6). Therefore, Hem 5 to Hem 9 were predicted to be granulocytes. These AMP-expressing clusters were broadly classified into two groups. One group was Hem 5, which expressed several AMPs, such as ALF-A1 and c-type lysozyme (Fig. 5 D and F), but also expressed other immune-related genes different from AMPs: *chitin-binding protein* and *virus responsible protein* (*VRP*) (Fig. 2). Chitin-binding protein is located on the cell surface and interacts with virus envelope proteins in shrimp (58). VRP is distributed in granulocytes, and infection by pathogenic viruses causes an increase in VRP transcripts (59). The other group is Hem 6 to 9, which strongly expresses major AMPs, such as penaeidin, stylicin, and SWD, suggesting that this group corresponds to the most studied immune-related cell subpopulation to date. The level of penaeidin-positive hemocytes is increased after bacterial infection (60), but is reduced after a virus infection (61). Stylicin and SWD mRNA expression was decreased after virus infection in *M. japonicus* and *P. monodon*.

**Figure 5.**
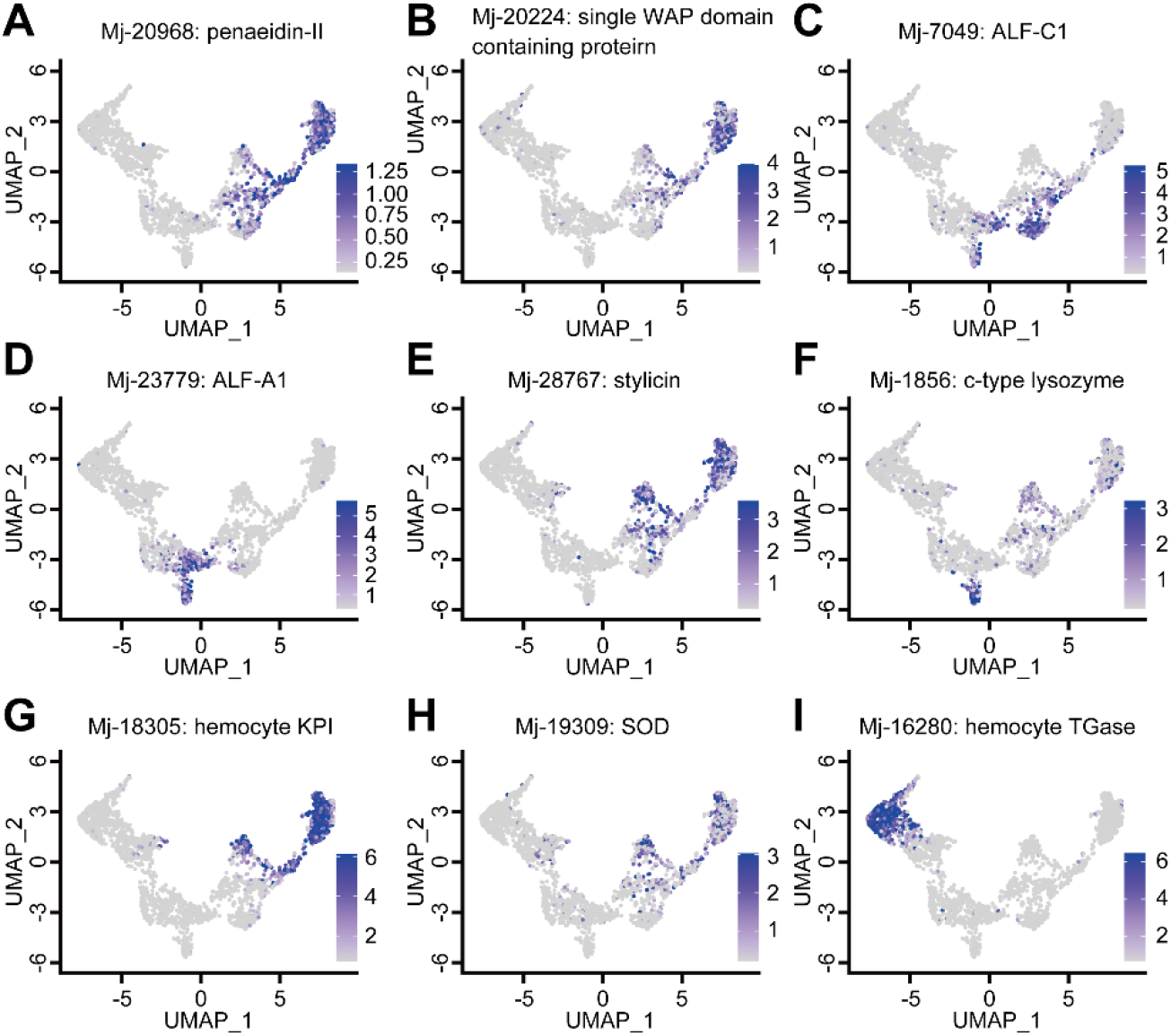
Expression pattern of the immune-related genes across cell clusters. (A) Penaeidin-II, (B) single WAP domain containing protein, (C) ALF-CA, (D) ALF-A1, (E) stylicin, (F) c-type lysozyme, (G) KPI, (H) SOD, and (I) TGase.

Together, the functions of these two groups can be characterized as follows. In Hem 5, VRP, whose level is known to be increased by viral infection, was specifically expressed, while AMPs were downregulated. Thus, Hem 5 is predicted to be the group that contributes to the immune response against viruses. Additionally, a virus infection causes an increase in the expression of both VEGF and its receptors in hemocytes to regulate a downstream signaling pathway (62-64). The high expression of PDGF/VEGF-related factors at Hem 5 (Fig. 2 and 4) supports our prediction that Hem 5 plays an essential role in shrimp biodefense against viruses. Previously, a subtype of granulocytes, termed semi-granular cells (SGC), were isolated from *L. vannamei*, in which lysozyme was highly expressed. It is possible that Hem 5 in *M. japonicus* corresponds to SGC found in *L. vannamei* because c-type lysozyme was strongly expressed in Hem 5, in our study. Conversely, the AMPs specifically expressed at Hem 6–9 were upregulated by bacterial infection. Therefore, these clusters contribute to bacterial defense. Different types of ALF play different roles in shrimp immunity, which improves the synergism in shrimp antimicrobial defenses (65). The expression patterns of ALFs were also different in each cluster (Fig. 5 C and D, Fig. S6 and Dataset S6), suggesting that different clusters have different immunological functions.

Some of the hemocyte-type-specific markers related to their immune function have been studied in crayfish: prophenoloxidase (proPO) in matured hemocytes, copper/zinc superoxide dismutase (SOD) in SGC, and Kazal-type proteinase inhibitor (KPI) in granular cells (GC), and transglutaminase (TGase) in immature cells (66). We BLAST searched homologies between these genes and common expressed genes to check their potential as specific markers. As a result, 31 genes were found to have homology with the genes listed above (Fig. S6 and Dataset S6). TGase was strongly expressed in immature clusters Hem 1 and Hem 2, and can, therefore, be used as a marker of immature hemocytes (Fig. 5 I and Fig. S6). ProPOs were not highly expressed in the present single-cell study. Both KPI and SOD were mostly expressed at both Hem 8 and Hem 9, and their expression levels were similar between these clusters (Fig. 5 G and H and Fig. S6). These results suggest that the functional segregation of hemocytes in shrimp is different from that in crayfish, in which these molecular markers are expressed distinctly between GC and SGC. The KPI of *M. japonicus* is expressed in only some hemocytes in healthy shrimps, and bacterial infection causes an increase in KPI expression (67). Therefore, it can be predicted that the number of KPI-positive hemocytes in Hem 8 and 9 would be increased upon a pathogen infection, in shrimp.

In the mosquito, scRNA-seq revealed a new subpopulation called “antimicrobial granulocytes” that expressed characteristic AMPs (22). Similarly, in *M. japonicus*, the expression patterns of immune-related genes were also different among certain clusters, suggesting that shrimp hemocytes are more heterogeneous than previously thought. It is anticipated that the class of granulocytes discussed in previous studies is actually a mixture of clusters exhibiting different roles.

### Validation of marker genes and the relationship between clusters and morphology

Our scRNA-seq results revealed nine major subpopulations and their marker genes, and the possible differentiation trajectory of *M. japonicus* hemocytes. Next, we examined the correlation between the morphology and expression of marker genes. Two major populations of hemocytes were sorted based on the forward versus side scatter plot obtained using microfluidic-based fluorescence-activated cell sorter (FACS). The sorted populations were observed using microscopy. FACS was able to separate hemocytes into two morphologically different populations (Fig. 6 A–E): smaller cells with low internal complexity in region 1 (R1) (50.7% ± 5.2%) and larger cells with high internal complexity in region 2 (R2) (47.7% ± 3.4%). From DIC and dye staining imaging (Fig. 6 A and B), we observed that cells in the R1 region contained no or few granules in the cytoplasm. The nucleus occupied a large portion of the volume in these cells (Fig. 6 C). Conversely, those in the R2 region had many granules in the cytoplasmic region, which occupied a large portion of cells (Fig. 6 D).

**Figure 6.**
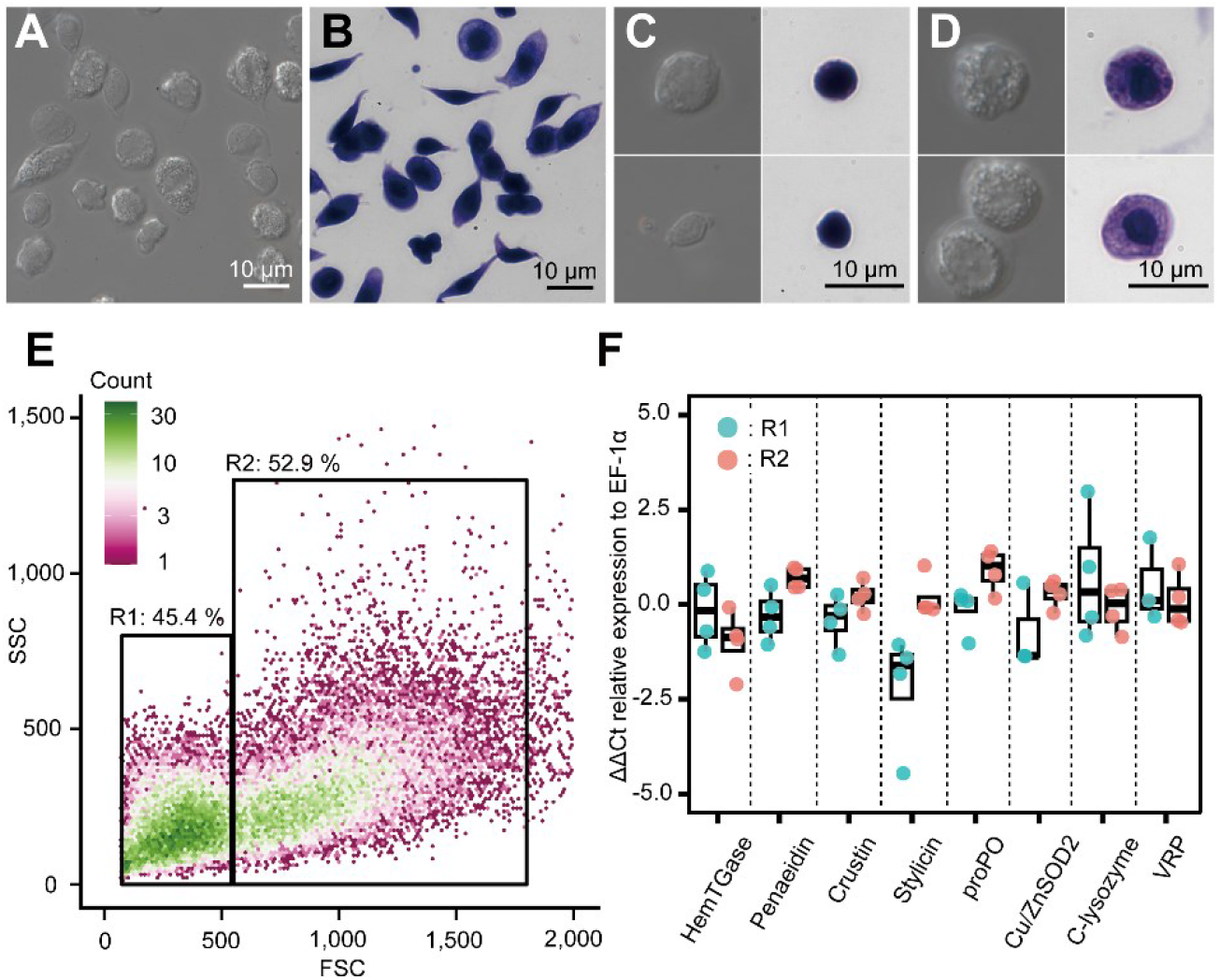
Morphological analysis of hemocytes and transcript profiles based on morphology. (A) DIC (Differential Interference Contrast) image of unsorted total hemocytes. (B) Dye stained total hemocytes. (C) DIC imaging and dye staining of R1 sorted hemocytes. (D) DIC imaging and dye staining of R2 sorted hemocytes. (E) FACS (fluorescence-activated cell sorting) analysis of hemocytes. Based on the FSC (forward scatter) and SSC (side scatter) two-dimensional space, two regions (R1 and R2) were obtained. (F) Differential gene expression analysis between R1 and R2 of hemocytes sorted using FACS. ΔΔCt values were analyzed using qRT-PCR. Higher ΔΔCt values indicate a higher accumulation of mRNA transcripts.

We conducted qRT-PCR analysis to determine the expression of some representative genes in these populations. The results showed that the ΔΔCt values of the transcripts of HemTGase were higher in R1 hemocytes than in R2 hemocytes, while the values of penaeidin, crustin, stylicin, proPO, and Cu/ZnSOD2 were higher in R2 hemocytes (Fig. 6 F). The ΔΔCt values of transcripts of c-type lysozyme and VRP were similar in R1 and R2 hemocytes. The combination of FACS and qRT-PCR results confirmed that the gene expression of the two populations in shrimp hemocytes was roughly divided based on morphology, that is, agranulocytes and granulocytes, which is consistent with our scRNA-seq analysis. Granulocytes found in the R2 region expressed major AMPs, such as penaeidin, crustin, and stylicin, suggesting that they consist of clusters Hem 6 to Hem 9. However, c-type lysozyme and VRP, which are markers of Hem 5 (Fig. 2), were expressed in cells in both R1 and R2 regions, indicating that Hem 5 exists in both populations and is indistinguishable from the morphological characteristics. The number of cells occupying R1 and R2 was 50.7% ± 5.2% and 47.7% ± 3.4% (from *n* = 4), while the total number of cells classified as Hem 1 – 4 and Hem 5 – 9 in our single-cell analysis was 50.4% (1,363 cells) and 49.6% (1,341 cells), respectively.

## Discussion

Our single-cell transcriptome analysis revealed that there are nine subpopulations of hemocytes in shrimp. This result differs from any previous classification strategy based on various approaches, such as simple staining (7-9), monoclonal antibodies (18), flow cytometry (68), and lectin-binding profiles (69). We hope that future studies will clarify the relationships between these phenotypic features and our classification to comprehend the role of each cluster in the immune system of shrimp.

The cluster-specific markers and cell proliferation-related genes found here helped us to understand how shrimp hemocytes differentiate. A strong expression of TGase, cell proliferation, and G2/M state-related genes in Hem 1 suggested that hemocytes in this cluster are oligopotent and located upstream in the differentiation process. In crustaceans, especially shrimp and crayfish, it is known that hemocytes are produced in HPT, and that differentiated hematopoietic cells from HPT circulate in the body fluid (9, 36, 66). However, G2/M state hemocytes of *M. japonicus* exist in the hemolymph and account for only 0.63% ± 0.28% of the circulating hemocytes (70). Hem 1 accounts for only 2.4% of the analyzed cells. This similarity in the fraction also indicates that only a very small fraction of oligopotent or initial state hemocytes exist among the circulating hemocytes.

Our results also revealed that the growth-related genes were expressed at specific clusters (Fig. 4 A and B). From these results, we propose that shrimp hemocytes differentiate as follows (Fig. 7 A and B): 1) Hem 1 is the initial state of circulating hemocytes and has an oligopotent ability, which leaked out from HPT; 2) Hem 3 is differentiated from Hem 1 by VWDE-like through Hem 2; 3) Hem 4 is differentiated from Hem 3 by TINAGL; 4) Hem 5 is differentiated from Hem 4 by PDGF/VEGF-related factor; 5) Hem 7 to Hem 9 are differentiated from Hem 4 by IGFBP-related protein through Hem 6; 6) CHF-like is expressed in Hem 9 to maintain the immature hemocytes, Hem 1 to Hem 4. Loss and gain function studies of these differentiating factors are necessary to prove the full differentiation process of penaeid shrimp hemocytes. Since we identified a group of proliferating cells and differentiation factors here, the next step is to establish a way to isolate and cultivate them. The markers of each cluster identified here will be good tools for isolating specific cell types.

**Figure 7.**
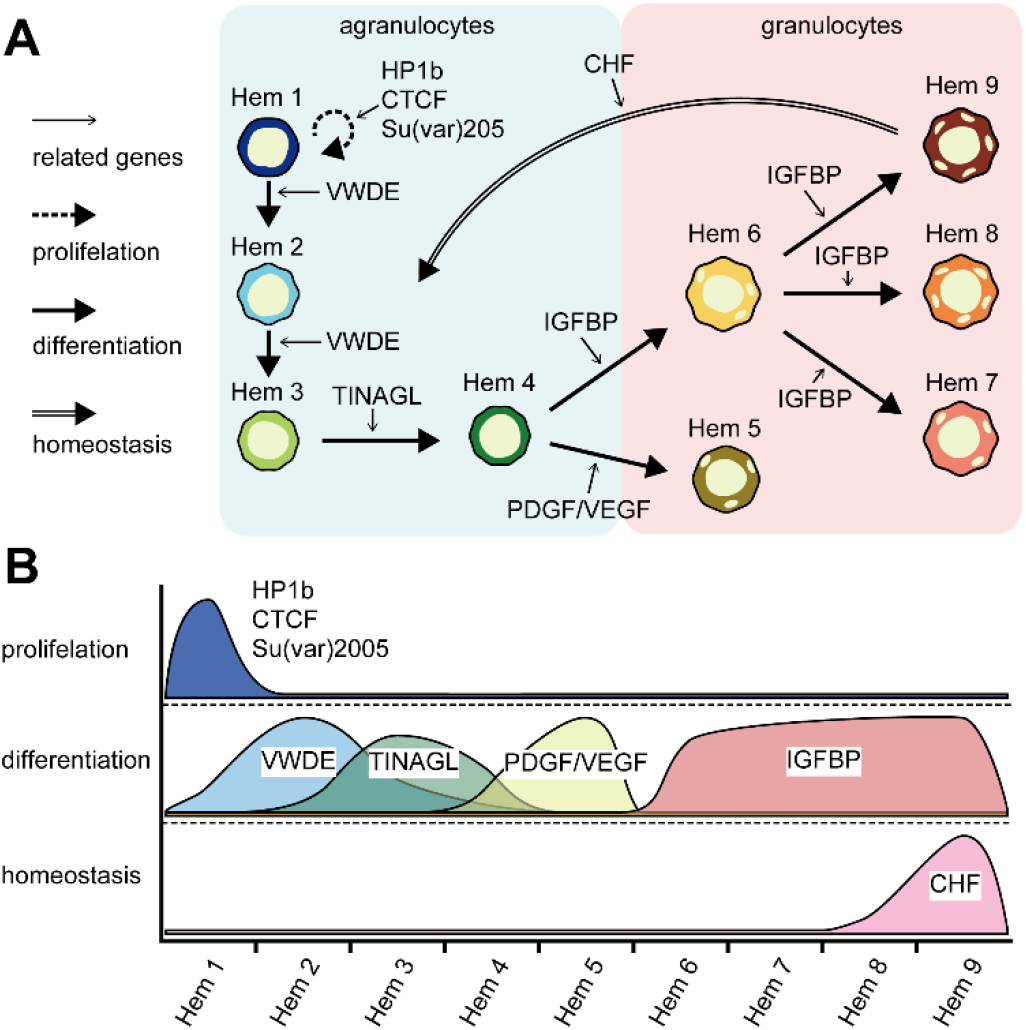
Model of the roles of cell growth-related genes in hemocyte differentiation. (A) Shrimp hemocyte differentiation process, 1) Hem 1 is the initial state of circulating hemocytes, which have oligopotent ability, which leaked out from the hematopoietic tissue (HPT); 2) Hem 3 is differentiated from Hem 1 by VWDE-like through Hem 2; 3) Hem 4 is differentiated from Hem 3 by TINAGL; 4) Hem 5 is differentiated from Hem 4 by PDGF/VEGF-related factor; 5) Hem 7 to Hem 9 are differentiated from Hem 4 by IGFBP-related protein through Hem 6; 6) CHF-like is expressed in Hem 9 to maintain the immature hemocytes, Hem 1 to Hem 4. (B) Schematics of cell growth or homeostasis-related gene expression across clusters. The gene expression distribution was cluster-specific.

Crayfish’s hemocytes and *Drosophila* hemocytes have been classified into three and four major types according to the marker genes, respectively (23, 36). However, these crayfish or *Drosophila* marker genes were found to be inadequate as cluster-specific markers for penaeid shrimp in our study. Insects and crustaceans are thought to become independent about 500 million years ago (71, 72), and shrimp and crayfish are thought to have become evolutionarily independent about 450 million years ago (73). Thus, it is straightforward to reason that the functions of these genes have changed during the evolutionary process. Therefore, we should not simply argue that the morphological and functional similarities between shrimp and *Drosophila*/crayfish hemocytes are the same.

Most of the markers that were characteristically expressed in Hem 1 and Hem 4 could not be functionally predicted by the BLAST search (Fig. 2). We speculate that the characteristic genes of Hem 1 are associated with cell duplication and that Hem 4 is associated with the regulation of cell differentiation. The division and differentiation mechanisms of shrimp hemocytes are still largely unknown, and no techniques on culture shrimp hemocytes *in vitro* have been reported. The analysis of these unknown gene characteristics may reveal these mechanisms. Notably, no specific marker gene was found in Hem 6, probably because these cells are yet to be specialized, unlike those in Hem 7 to Hem 9.

In conclusion, we succeeded in classifying shrimp hemocytes into nine subpopulations based on their transcriptional profiles, while they were only classified into two groups using FACS. Furthermore, our results imply that hemocytes differentiate from a single initial population. Although we have not yet successfully cultured crustacean hemocytes in passaging cultures, information on these subpopulations and marker genes will provide a foothold for hemocyte culture studies. Despite our success in the classification of hemocytes, we have not yet been able to fully understand the functions of each hemocyte group in detail. One reason for this is that the functions of some marker genes are still unknown. The present single-cell transcriptome data serves as a platform providing the necessary information for the continuous study of shrimp genes and their functions. Additionally, we have only determined the subpopulations of hemocytes in the normal state, so that our next goal will be to analyze the hemocytes in the infected state of certain diseases at different levels of cell maturation. In this way, we will be able to identify the major subpopulations that may work against the infectious agent. Likewise, it will be interesting to see how hemocytes from fertilized eggs mature. Additionally, single-cell analysis of hematopoietic tissues can also be expected to reveal more detailed differentiation mechanisms. Unlike terrestrial invertebrates, such as *Drosophila* and mosquitoes, shrimps are creatures that live in the ocean and are evolutionarily distant from each other. Understanding the immune system of shrimp species will require continuous effort, but we believe that it will provide efficient solutions to aquaculture problems.

## Materials and Methods

### Shrimp and cell preparation

Twenty-three female kuruma shrimp, *Marsupenaeus japonicus*, with an average weight of 20 g, were purchased from a local distributor and maintained in artificial seawater with a 34 ppt salinity with a recirculating system at 20°C. Hemolymph was collected using an anticoagulant solution suitable for penaeid shrimp (7) from an abdominal site. The collected hemolymph was centrifuged at 800 x *g* for 10 min to collect the hemocytes, which were then washed twice with PBS and the osmolarity was adjusted to kuruma shrimp (KPBS: 480 mM NaCl, 2.7 mM KCl, 8.1 mM Na_2_HPO_4_·12H_2_O, 1.47 mM KH_2_PO_4_, pH 7.4).

### Preparation of expressing gene list of hemocytes

*De novo* assembled transcript data were prepared as a reference for mapping Drop-seq data, because the genome sequence of *M. japonicus* is still unknown. To improve the quality of the *de novo* assembled transcript sequences, we prepared long-read mRNA sequences using MinION (Oxford Nanopore Technologies) direct RNA sequencing to conduct hybrid *de novo* assembly. Poly(A) tailed RNA was purified from 58 μg of total RNA from the hemocytes of 16 shrimp using Dynabeads Oligo(dT)_25_ (Thermo Fisher Scientific), and 500 ng of poly-(A) RNA was ligated to adaptors using a direct RNA sequencing kit (Oxford Nanopore Technologies) according to the manufacturer’s manual version DRS_9080_v2_revL_14Aug2019. Finally, 44 ng of the library was obtained and sequenced using MinION, by using a MinION flow cell R9.4.1 (Oxford Nanopore Technologies). All sequencing experiments were performed using MinKNOW v3.6.5 without base calling. Raw sequence data were then base-called using Guppy v3.6.1. Once the raw signal from the MinION fast5 files was converted into fastq files, the sequencing errors were corrected using TALC v1.01 (74)(https://gitlab.igh.cnrs.fr/lbroseus/TALC) by using the Illumina short reads sequence of kuruma shrimp hemocytes (DDBJ Sequence Read Archive (DRA) accession number DRA004781). The corrected long-read sequences from MinION and short-read sequences from Illumina Miseq were hybrid *de novo* assembled using rnaSPAdes v3.14.1 (75)(https://cab.spbu.ru/software/rnaspades/) and Trinity 2.10.0 (76)(https://github.com/trinityrnaseq/trinityrnaseq/wiki). All assembled *de novo* transcripts were merged and subjected to the EvidentialGene program v2022.01.20 (http://arthropods.eugenes.org/EvidentialGene/) to remove similar sequences with a default parameter. The remaining sequences were renamed as Mj-XXX and used as a hemocyte-expressing gene list. The assembled sequences and code used to perform base-calling and *de novo* assembly are available on GitHub at https://github.com/KeiichiroKOIWAI/Drop-seq_on_shrimp.

### Single-cell and single-bead encapsulation by a microfluidic device and exonuclease and reverse transcribe reaction on a bead

The drop-seq procedure was used to encapsulate single hemocytes and single mRNA capture beads together into fL-scale microdroplets, as previously described (19). The following steps were performed in triplicates for three shrimp individuals. Briefly, the self-built Drop-seq microfluidic device was prepared by molding polydimethylsiloxane (PDMS; Sylgard 184, Dow Corning Corp.) from the microchannel structure formed by the negative photoresist (SU-8 3050, Nippon Kayaku Co.). Using this device, droplets containing a cell and a Barcoded Bead SeqB (ChemGenes Corporation) were produced up to 2 mL per sample using a pressure pump system (Droplet generator, On-chip Biotechnologies Co., Ltd.). During the sample introduction, the vial bottles containing cells and beads were shaken using a vortex mixer to prevent sedimentation and aggregation (77). Droplets were collected from the channel outlet into the 50-mL corning tube and incubated at 80°C for 10 min in a water bath to promote hybridization of the poly(A) tail of mRNA and oligo d(T) on beads. After incubation, droplets were broken promptly and barcoded beads with captured transcriptomes were reverse transcribed using Maxima H Minus Reverse Transcriptase (Thermo Fisher Scientific) at RT for 30 min, then at 42°C for 90 min. Then, the beads were treated with Exonuclease I (New England Biolabs) to obtain single-cell transcriptomes attached to microparticles (STAMP). The first-strand cDNAs on beads were amplified using PCR. The beads obtained above were distributed throughout PCR tubes (1,000 beads per tube), wherein 1× KAPA HiFi HS Ready Mix (KAPA Biosystems) and 0.8 µM 1^st^ PCR primer were included in a 25 μL reaction volume. PCR amplification was achieved using the following program: initial denaturation at 95°C for 3 min; 4 cycles at 98°C for 20 s, 65°C for 45 s, and 72°C for 6 min; 12 cycles of 98°C for 20 s, 67°C for 20 s, and 72°C for 6 min; and a final extension at 72°C for 5 min. The amplicons were pooled, double-purified with 0.9× AMPure XP beads (Beckman Coulter), and eluted in 100 μL of ddH_2_O. Sequence-ready libraries were prepared according to Picelli, Bjorklund (78). A total of 1 ng of each cDNA library was fragmented using home-made Tn5 transposome in a solution containing 10 mM TAPS-NaOH (pH 8.5), 5 mM MgCl_2_, and 10% dimethylformamide at 55°C for 10 min. The cDNA fragments were purified using a DNA Clean & Concentrator Kit (Zymo Research) and eluted in 25 µL of ddH_2_O. The index PCR reaction was performed by adding 12 µL of the elute to a mixture consisting of 1× Fidelity Buffer, 0.3 mM dNTPs, 0.5 U KAPA HiFi DNA polymerase (KAPA Biosystems), 0.2 µM P5 universal primer, and 0.2 µM i7 index primer. Each reaction was achieved as follows: initial extension and subsequent denaturation at 72°C for 3 min and 98°C for 30 s; 12 cycles of 98°C for 10 s, 63°C for 30 s, and 72°C for 30 s; and a final extension at 72°C for 5 min. The amplified library was purified using 0.9× AMPure XP beads and sequenced (paired-end) on an Illumina NextSeq 500 sequencer (NextSeq 500/550 High Output v2 kit (75 cycles); 20 cycles for read1 with custom sequence primer, 8 cycles for index read, 64 cycles for read2. Before performing the Drop-seq on kuruma shrimp hemocytes, we validated the protocol by performing the same procedure using a mixture of HEK293 and NIH3T3 cells and sequencing the test library using a Miseq Reagent Kit v3 (150 cycles).

### Analysis of single-cell data

Paired-end reads were processed and mapped to the reference *de novo* assembled gene list of hemocytes following the Drop-seq Core Computational Protocol version 2.0.0, and the corresponding Drop-seq tools v2.3.0 (https://github.com/broadinstitute/Drop-seq) provided by McCarroll Lab (http://mccarrolllab.org/dropseq/). The Picard suite (https://github.com/broadinstitute/picard) was used to generate the unaligned bam files. The steps included the detection of barcode and UMI sequences, filtration and trimming of low-quality bases and adaptors or poly(A) tails, and the alignment of reads using bowtie2 v2.4.1 (79) (http://bowtie-bio.sourceforge.net/bowtie2/index.shtml). The cumulative distribution of reads from the aligned bam files was obtained using BAMTagHistogram, and the number of cells was inferred using Drop-seq tools.

### Data integration

After digital expression data from 3 shrimps were read using Seurat v3.2.1 (80, 81)(https://satijalab.org/seurat/), SCTransform (81) was performed to remove the technical variation and to select common expressing genes, while retaining biological heterogeneity. We ran a PCA using the expression matrix of the top 3,000 most variable genes. The total number of principal components (PCs) required to compute and store was 50. The UMAP was then performed using the following parameters: n.neighbors, min.dist, and n.components were 10L, 0.1, and 2, respectively, to visualize the data in the two-dimensional space, and then the clusters were predicted with a resolution of 0.5.

### Functional prediction of commonly expressed genes

The predicted functions of selected assembled sequences as common expressing genes across three replicates were searched using BLAST program v2.2.31 (82, 83) (https://ftp.ncbi.nlm.nih.gov/blast/executables/blast+/LATEST/) on penaeid shrimp identical proteins (downloaded from NCBI Identical Protein Groups on 19th of August, 2020) with the blastx parameter of e-value as 0.0001 and num_alignments as 3. Then, the functions of each cluster were predicted based on the marker genes. Marker genes were predicted using the Seurat FindAllMarkers tool with the following parameters: min.pct as 0.5, logfc.threshold as 1, and test.

### Visualization of genes with distinctive functions on single hemocytes

To visualize the genes with distinctive functions of shrimp, we extracted the distinct sequences from commonly expressed genes based on their blastx results against penaeid shrimp identical proteins. Here, we focused on the cell growth-related genes listed in Dataset S5 and on immune-related genes, such as antimicrobial peptides (AMPs), transglutaminase, copper/zinc superoxide dismutase, and Kazal-type proteinase inhibitors listed in Dataset S6. From these genes that showed a characteristic expression among single-cell data, dot plots, violin plots, or feature plot visualizations were applied using Seurat functions.

### Comparison with *Drosophila* marker genes

To check whether the *Drosophila* marker genes are applicable to shrimp, we performed a BLAST search on *the Drosophila* cell cycle and cell type markers (https://github.com/hbc/tinyatlas). Common genes among the three shrimp species were tblastx searched for *Drosophila melanogaster* genes (dmel-all-gene-r6.34.fasta; downloaded from FlyBase https://flybase.org/) with the parameters of e-value as 0.0001 and num_alignments as 3.

### Pseudo temporal ordering of cells using Monocle 3

The integrated data of Seurat were transferred to Monocle 3 (84) (https://github.com/cole-trapnell-lab/monocle3) to calculate a cell trajectory using the learn_graph function. We assigned the start point based on the expression of cell proliferation-related genes and *Drosophila* marker genes of the cell cycle.

### Cell sorting of hemocytes and qRT-PCR of marker genes

To validate the Drop-seq results on hemocytes, populations of hemocytes in the forward scatter (FSC) and side scatter (SSC) two-dimensional space were sorted using a microfluidic cell sorter (On-chip sort, On-chip Biotechnologies Co., Ltd.) from four shrimp individuals (Fig. S7). In the FSC/SSC two-dimensional space, two main populations were predicted as Region1 (R1): small/simple and Region2 (R2): large/complexity populations, which were defined as agranulocytes and granulocytes, respectively. After sorting, some sorted hemocytes were immediately fixed in 2% formalin in KPBS and stained with a May-Grunwald and Giemsa staining solution to observe the cellular components. Non-stained and stained hemocytes were subjected to microscopy IX71 (Olympus Corporation) to observe their structures.

Total RNA was also collected from sorted cells and pre-sorted cells. The concentration of RNA was measured using a nanodrop, and cDNA was transcribed using a High-Capacity cDNA Reverse Transcription Kit (Thermo Fisher Scientific). Constructed cDNA was diluted five times with TE buffer and subjected to qRT-PCR using KOD SYBR qPCR (TOYOBO Co. Ltd.), following the manufacturer’s protocol. The expression of each gene was calculated using the ΔΔCT method against elongation factor-1 alpha and total hemocytes.

### Data files and analysis code

The raw sequence data of newly sequenced *M. japonicus* transcriptomic reads were archived in the DDBJ Sequence Read Archive (DRA) of the DNA Data Bank of Japan as follows: MinION mRNA direct sequencing: DRA010948; Drop-seq shrimp rep1: DRA010950; shrimp rep2: DRA010951; shrimp rep3: DRA010952; mixture sample of HEK293 and 3T3 cells: DRA010949. The Seurat digital expression data were archived in the Genomic Expression Archive of the DNA Data Bank of Japan: E-GEAD-403. Fast5 data of MinION direct RNA sequencing will be made available upon request from the authors. The code used to perform *de novo* assembly, clustering, and marker analysis is available on GitHub at https://github.com/KeiichiroKOIWAI/Drop-seq+on+shrimp. The key resources are listed in Dataset S7.

## Supporting information

Supplementary Fig S1 to S7 and Legends for Dataset S1 to S7

## Acknowledgments

We would like to thank Fumiko Sunaga for her technical support in performing cell sorting and analysis, and Editage (www.editage.com) for English language editing. This work was supported by JSPS KAKENHI (JP19J00539 and JP20K15603) to K. Koiwai; a Grant-in-Aid for Scientific Research on Innovative Areas (17H06425) to K. Kikuchi.

