## Supplementary Fig S1 to S7 and Legends for Dataset S1 to S7 for "Single-cell RNA-seq analysis reveals penaeid shrimp hemocyte subpopulations and cell differentiation process"

This PDF file includes:

Supplementary Fig S1 to S7

Legends for Dataset S1 to S7

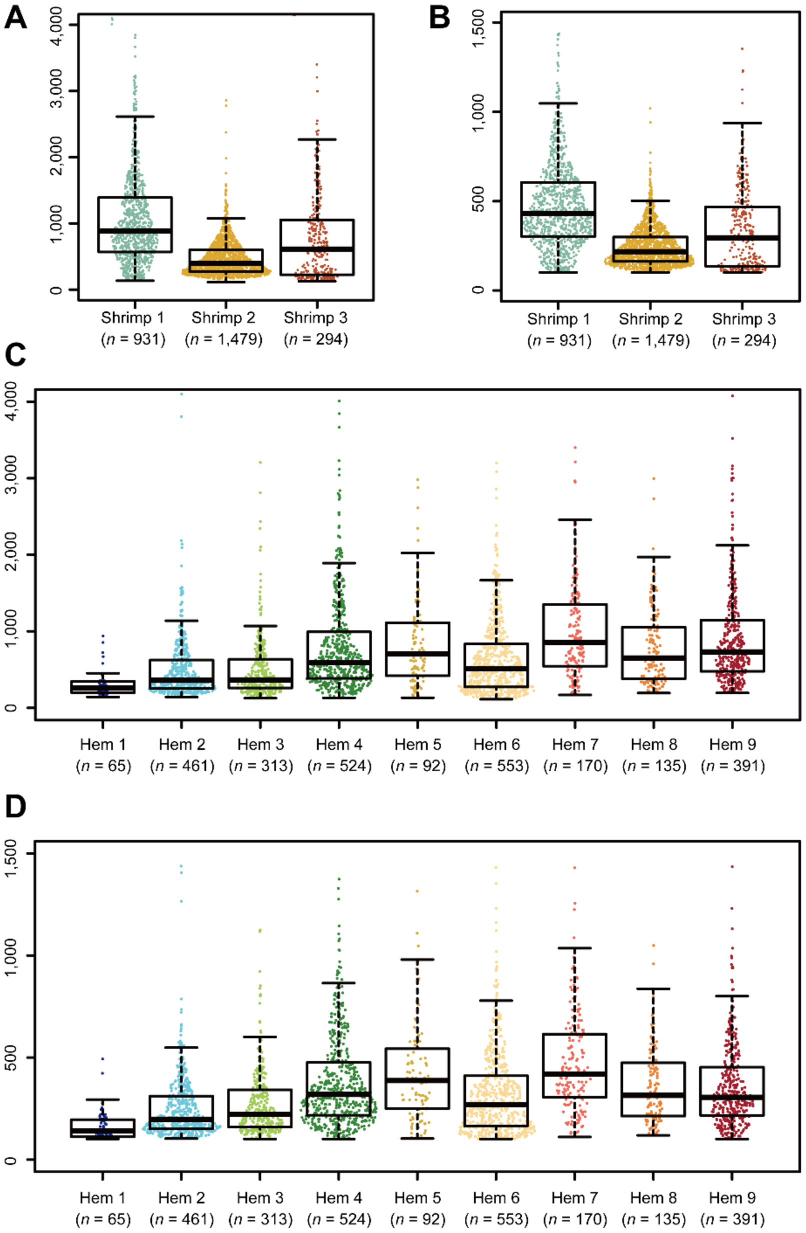


**Fig. S1.** Violin plots show the distribution of the number of transcripts (scored by UMIs) (A) and genes (B) detected per hemocyte on individual shrimp. Violin plots show the distribution of the number of transcripts (scored by UMIs) (C) and genes (D) detected per hemocyte on each cluster.

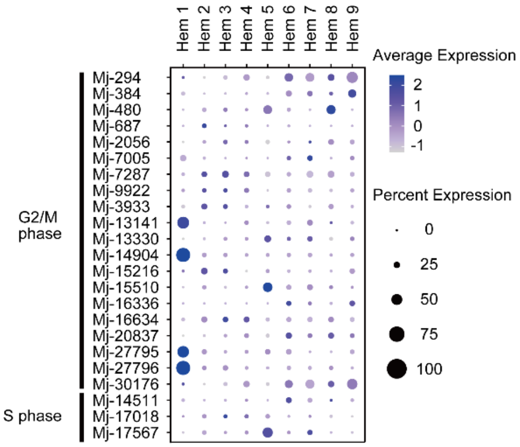


**Fig. S2.** Dot plot representing the *Drosophila* cell cycle markers per cluster based on average expression. Color gradient of the dot represents the expression level, while the size represents the percentage of cells expressing any gene per cluster.

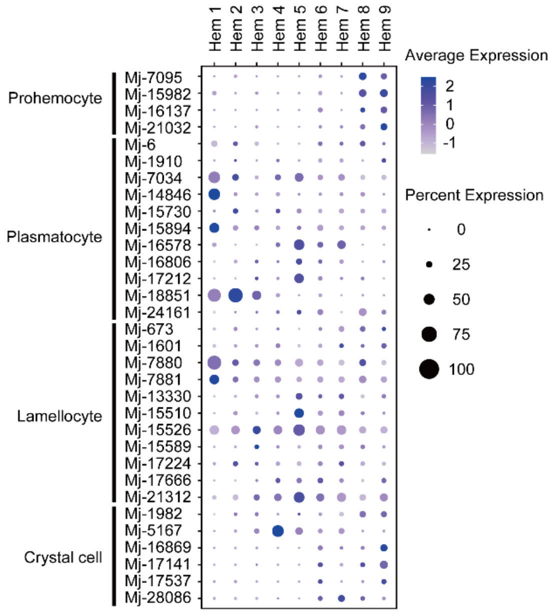


**Fig. S3.** Dot plot representing the *Drosophila* hemocyte type markers per cluster based on average expression. Color gradient of the dot represents the expression level, while the size represents the percentage of cells expressing any gene per cluster.

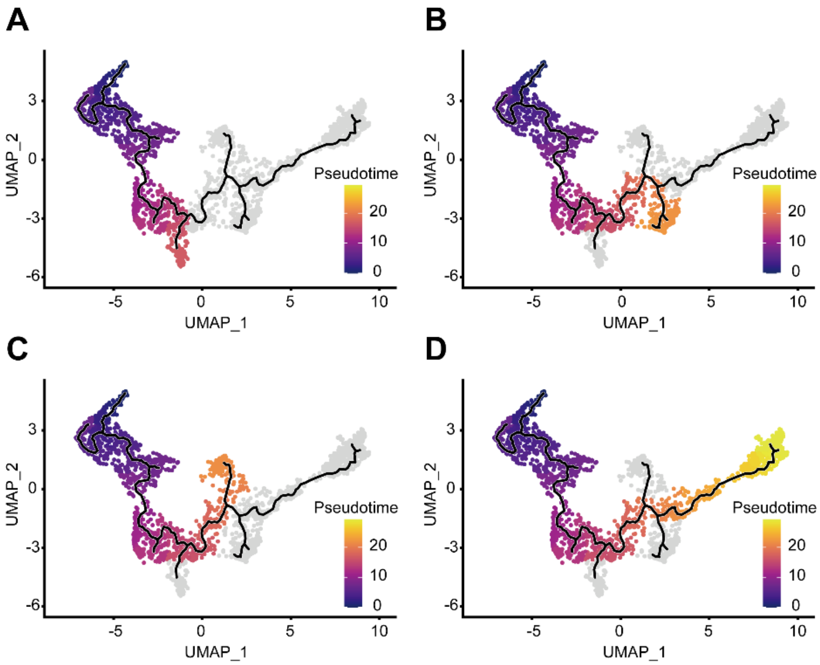


**Fig. S4.** Four major lineage pathways of hemocytes were obtained from the start site: Lineage 1 (A), Lineage 2 (B), Lineage 3 (C) and Lineage 4 (D).

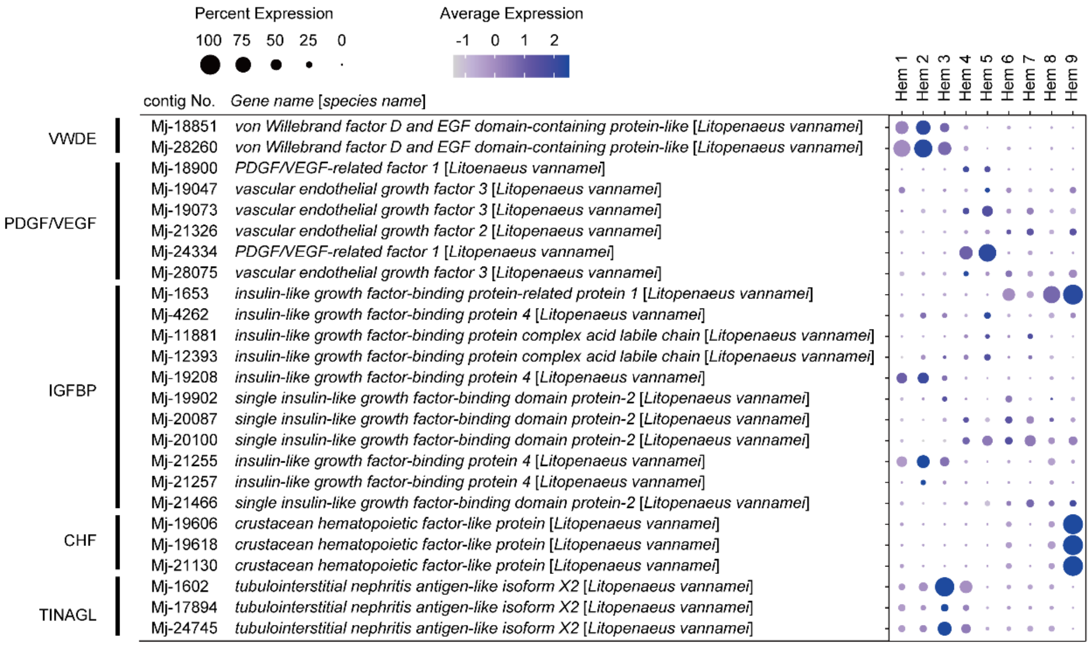


**Fig. S5.** Dot plot representing the cell growth related genes per cluster based on average expression. Color gradient of the dot represents the expression level, while the size represents the percentage of cells expressing any gene per cluster.

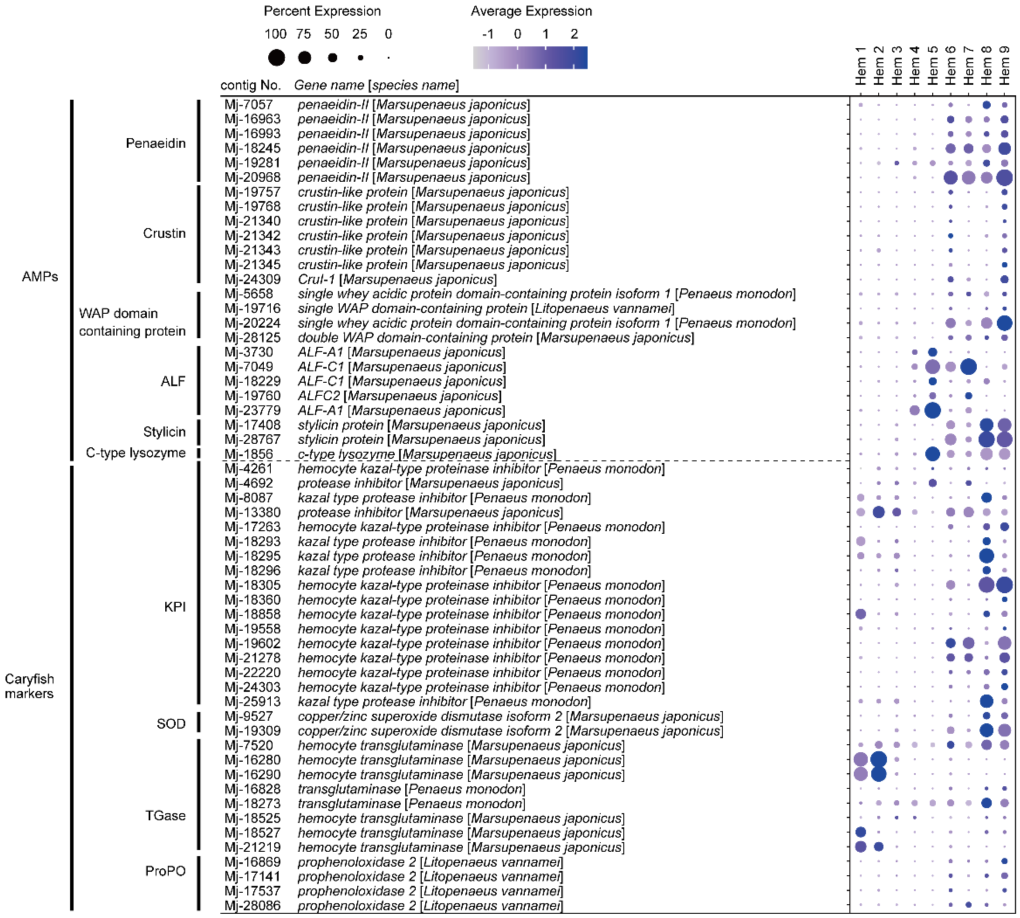


**Fig. S6.** Dot plot representing the immune related genes per cluster based on average expression. Color gradient of the dot represents the expression level, while the size represents the percentage of cells expressing any gene per cluster.

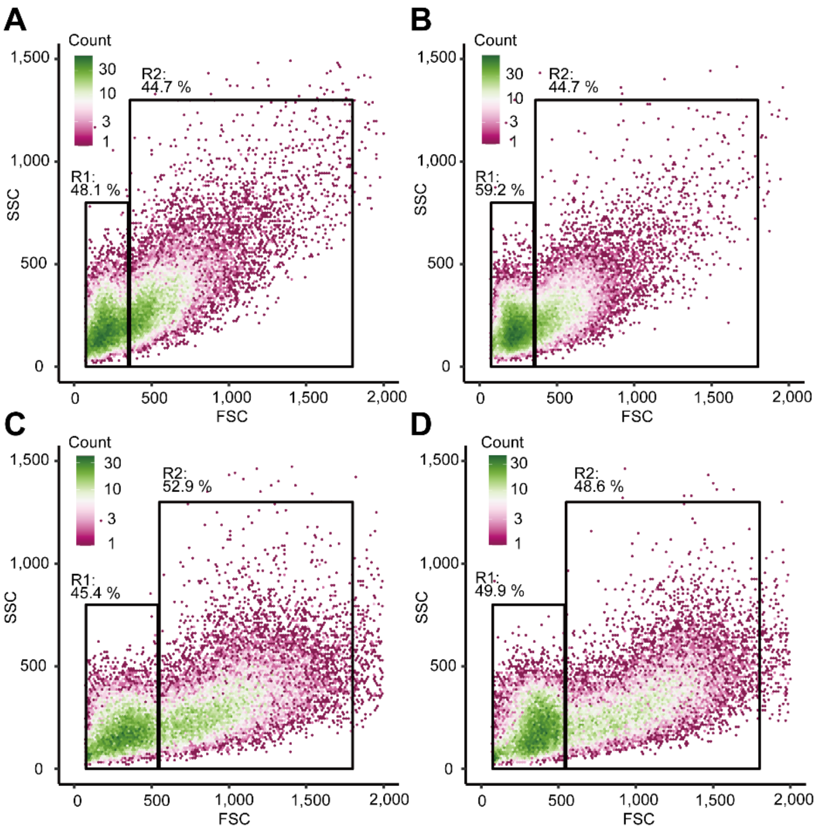


**Fig. S7.** FACS (fluorescence-activated cell sorting) analysis of hemocytes from four individual shrimps (A-D). Based on the FSC (forward scatter) and SSC (side scatter) two-dimensional space, two regions (R1 and R2) were obtained.

Legends for Dataset S1 to S7

Dataset S1. Blastx researching of assembled genes against penaeid shrimp's identical proteins.

Dataset S2. Predicted markers on each cluster.

Dataset S3. Cell cycle associate genes predicted by blast searching of assembled genes against *Drosophila* melanogaster's genes.

Dataset S4. Cell type marker genes predicted by blast searching of assembled genes against *Drosophila* melanogaster's genes.

Dataset S5. Cell growth related genes from assembled genes.

Dataset S6. Anti-microbial peptides and crayfish hemocyte marker genes from assembled genes.

Dataset S7. Key resource table
